# Scaling protein-water interactions in the Martini 3 coarse-grained force field to simulate transmembrane helix dimers in different lipid environments

**DOI:** 10.1101/2022.09.09.506752

**Authors:** Ainara Claveras Cabezudo, Christina Athanasiou, Alexandros Tsengenes, Rebecca C. Wade

## Abstract

Martini 3, the latest version of the widely used Martini force field for coarse-grained molecular dynamics simulations, is a promising tool to investigate proteins in phospholipid bilayers. However, simulating other lipid environments, such as detergent micelles, presents challenges due to the absence of validated parameters for their constituent molecules. Here, we propose parameters for the micelle-forming surfactant, dodecylphosphocholine (DPC). These result in micelle assembly with aggregation numbers in agreement with experimental values. However, we identified a lack of hydrophobic interactions between transmembrane helix protein dimers and the tails of DPC molecules, preventing insertion and stabilization of the protein in the micelles. This problem was also observed for protein insertion by self-assembling 1-palmitoyl-2-oleoyl-sn-glycero-3-phosphocholine (POPC) or dipalmitoylphosphatidylcholine (DPPC) bilayers. We propose the reduction of the non-bonded interactions between protein and water beads by 10% as a simple and effective solution to this problem that enables protein encapsulation in phospholipid micelles and bilayers without altering protein dimerization or bilayer structure.

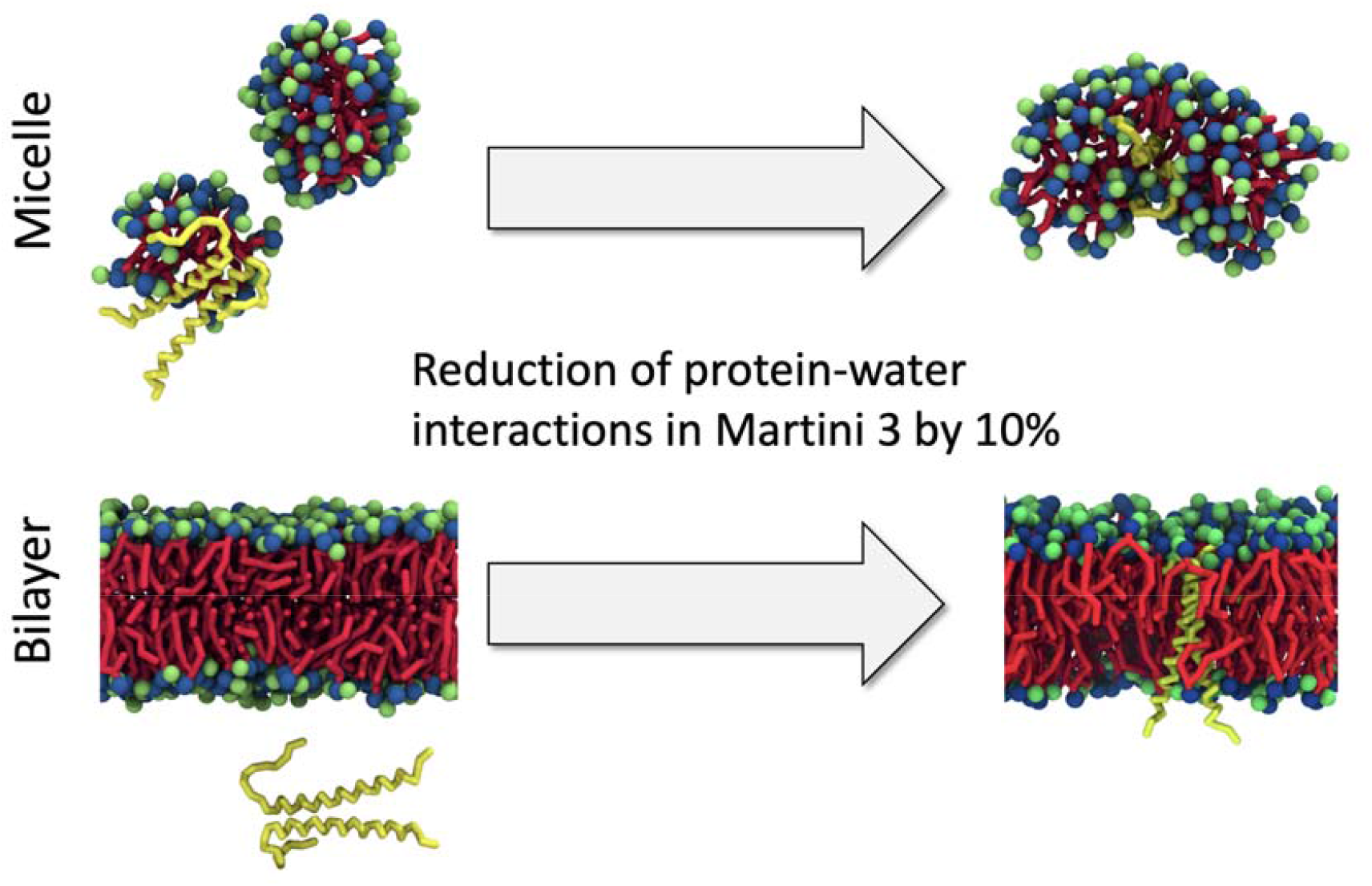

## INTRODUCTION

Molecular dynamics (MD) simulation is a powerful approach to explore the behaviour of biomolecules at a spatio-temporal resolution that cannot be obtained with experimental techniques.^1 2^ This method can, however, be limited by a lack of sampling of the relevant configuration space, which thus compromises the plausibility of the results obtained. In atomistic MD simulations, the length of the simulations is limited by the high number of computationally expensive calculations required. Coarse-grained (CG) simulations alleviate this problem by grouping several atoms into “beads”, thus reducing the number of degrees of freedom, and therefore, the number of calculations required. Thus, for a given amount of computation time, much longer simulation times can be achieved.

The Martini force field is one of the most widely used CG models due to its good performance and simplicity. Contrary to all-atom calculations, this method does not aim to provide a detailed structural view of the system, but to provide a broader picture of the processes taking place over a longer time scale.^3^ Martini beads usually represent groups of four non-hydrogen atoms and have different degrees of polarity. Parameters for specific beads are assigned against thermodynamic data, particularly oil/water partition coefficients. The Martini model has been shown to be able to reproduce a considerable number of experimental properties.^4^

A new, improved version of the Martini force field has recently been released.^5^ This improved version, referred to as Martini 3, overcomes some problems that have been reported for previous versions of the force field, such as an excessive aggregation of soluble and transmembrane proteins,^6 7^ or an excessive stacking of nucleotides.^8 9^ One of the most important changes in Martini 3 is the introduction of a specific type of bead for water (W), which has been parameterized separately from the rest of the beads.

Since its release, Martini 3 has been reported to present a number of issues, such as numeric instability of simulations with large time steps and an underestimation of the dimensions of intrinsically disordered proteins (IDPs).^10 11^ An increase of the bead size or the mass of ion beads has been proposed to enable simulations with large time steps, whereas an up-scaling of non-bonded interactions between protein and water beads resolved the IDP problem.

Detergent micelles are frequently used to experimentally determine the structure of transmembrane proteins, as they mimic the hydrophobic environment of phospholipid bilayers.^12 13^ Hence, many experimental structures obtained in micelles, are used as starting structures in molecular dynamics simulations. The fact that these simulations are usually run for the proteins in phospholipid bilayers hinders a direct comparison to experimental data. The simulation of these transmembrane proteins in detergent micelles is thus necessary to understand the influence of the lipid environment on their three-dimensional structure and dynamics. However, parameters for micelle-forming lipids in Martini 3 are currently not available. To overcome this problem, the widely used surfactant dodecylphosphocholine (DPC) was parameterized in this study on the basis of the existing parameters for other phospholipids in Martini 3 and using the parameters of DPC in the previous version of the force field, Martini 2, as a reference. Simulations at different lipid concentrations showed that the newly parameterized DPC molecules were able to assemble into micelles with properties similar to those obtained by different experimental techniques. However, we found that transmembrane (TM) helix dimer proteins fail to insert into the DPC micelles in Martini 3: they interacted with water beads or with the polar head groups of surfactant molecules rather than with their hydrophobic tails. Moreover, the transmembrane helix proteins rarely inserted into 1-palmitoyl-2-oleoyl-sn-glycero-3-phosphocholine (POPC) or dipalmitoylphosphatidylcholine (DPPC) bilayers during bilayer self-assembly. We here show that reducing protein-water interactions by introducing a small uniform scaling factor of the Lennard-Jones interactions in Martini 3 overcomes this aberrant behaviour for TM helix dimer proteins, leading to their correct insertion in micelles and self-assembled phospholipid bilayers.

## MATERIAL AND METHODS

### System preparation

The structure of the TM helix dimer of glycophorin A (GpA), which was obtained by NMR in DPC micelles, was used in most of the simulations in which protein was present (PDB ID: 1AFO^14^). The martinize2.py script ^5^ was used with DSSP^15^ to map atomistic structures to CG models. An elastic network was applied to the helical region of the TM monomers (residues 73 to 95) using a force constant of 700 kJ/mol nm² and distance cut-offs of 0.5 and 0.9 nm to preserve secondary structure, while retaining the intrinsic flexibility of the N-loop and C-loop regions.

Some test simulations were also run for the TM helix dimer of the tyrosine kinase neurotrophin receptor TrkA, whose structure was also determined by NMR in DPC micelles (PDB ID: 2N90^16^). The CG model of TrkA was obtained from the atomistic structure using an analogous procedure to that for GpA.

To explore micelle formation at different surfactant concentrations, simulations were run in boxes of different sizes and with different numbers of DPC molecules. In all cases, DPC molecules were placed at random positions at the beginning of the trajectory. Each system was solvated in water and neutralized in 0.15 M NaCl, using the standard Martini 3 ion parameters.

To investigate bilayer self-assembly, 250 POPC or DPPC molecules and the protein dimer were placed at random positions in a 10&10&10 nm^3^ with subsequent solvation in water and neutralization in 0.15 M NaCl. Some simulations were also initiated using the *INSANE* method to insert proteins into a preformed POPC bilayer.^17^

### CG-MD simulations

CG simulations were performed using the 2020.4 version of the GROMACS software package^18^ and the Martini 3 force field.^5^ A series of parameters which have been previously shown to boost performance were used.^19^ These include the use of the Verlet neighbour search (VNS) algorithm and cut-offs of 1.1 nm for Lennard Jones and Coulomb potentials. Long range electrostatic interactions were treated using the reaction-field method with a dielectric constant of 15. The velocity rescale thermostat and the Parrinello-Rahman barostat ^20 21^ were used with coupling parameters of 1.0 and 12.0 ps, respectively. A compressibility of 3e^-4^ was used, as well as a temperature of 310 K and a pressure of 1 bar. A time step of 20 fs was set and the neighbour list was updated every 10 time steps. Trajectory frames were saved for analysis every 1 ns.

For each system, different replicas were run using random initial velocities and positions (A summary of all simulations can be found in **Supplementary Table 1**). After an energy minimization step, equilibration and production runs were performed in the NPT ensemble. Position restraints with a force constant of 4000 kJ/mol nm² were applied to the backbone beads of the TM region of the protein during equilibration and released for production. Simulations of DPC micelles were run using isotropic pressure coupling. In simulations of bilayer self-assembly, isotropic pressure coupling was used for the first 60 ns of equilibration. Bilayer formation was visually assessed afterwards, and, if required, a rotation of the entire system was applied to place the bilayer in the X-Y plane. The remainder of the simulation was then run using semi-isotropic pressure coupling.

Some simulations were run with Martini 2.2 in order to compare the two versions of the force field. In these Martini 2.2 simulations, cut-offs of 1.2 nm were used for the Lennard-Jones and Coulomb potentials. The non-polarizable water model was used in the Martini 2.2 simulations. All other simulation parameters were the same as for the Martini 3 simulations. All simulations in bilayers were equilibrated for 2 *µ*s, which were not included in the analysis. Due to the long equilibration protocol, all production frames were used for analysis of the trajectories. To assess micelle and bilayer self-assembly, the equilibration was considered for analysis, as lipid arrangement takes place during this step.

### Scaling of non-bonded interactions in Martini 3

In order to scale the non-bonded interactions of the force field, a uniform scaling factor was applied on energy well depths *ε*_ij_ of the Lennard Jones potentials of either all protein-lipid hydrophobic C1 bead pairs or all protein-water bead pairs, as has been described previously.^22^

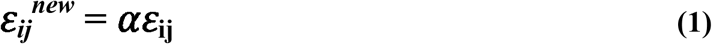

### Data analysis and visualisation

Some basic analysis tasks, such as the calculation of the root mean square deviation (RMSD) or temperature and pressure monitoring were performed using the tools provided in the GROMACS software package. More complex analyses, such as the calculation of dihedral angles or the distances between centers of geometry of the two helices were implemented in the Python programming language ^23^ with extensive use of the MDAnalysis ^24 25^, NumPy ^26^ and Pandas ^27^ packages.

In order to evaluate the exploration of the conformational landscape, kernel density heatmaps for different structural parameters were plotted using the Seaborn Python package.^28^ Further data analysis was performed using the R language for statistical computing. ^29^ PyMol^30^ and VMD ^31^ were used for visualization of the trajectories, and the Tachyon ray-tracer was used for image generation.^32^

### Structural parameters of transmembrane helix dimers

To explore the conformational space of the helix dimers, three different dihedral angles were defined. These were calculated after centering on one of the two helices (referred to as the “first”) and aligning each frame of the trajectory to this centred helix. The *crossing angle* was defined as the dihedral angle between the vectors that connect the centroid of each helix with the centroid of its upper half.^33^ Furthermore, rotation angles of the second helix with respect to the first helix and to itself, referred to as *position* and *phase*, respectively, were calculated. The position is represented by the dihedral angle between the vector connecting the centroid of each helix and the vector that connects the centroid of the upper half of the first helix with a residue located in the dimer interface in the same helix.^34^ To calculate the phase, the dihedral angle between the line that passes through the centroids of both helices and the one that connects the reference residue in the second helix and the centroid of its upper half was computed.

### Contact analysis

To compute the percentage occupancies of contacts between TM helix dimers during the simulations, all replicas were merged into a single trajectory and the whole system was aligned to one TM helix. Broken molecules (due to the use of a periodic simulation box) were made whole using the *gmx trjconv* command in GROMACS. By TCL scripting, contacts between the beads of the two helices were defined using a cut-off distance of 6 Å and contact occupancies were calculated for each bead pair. The contact occupation of a pair of residues was defined as the highest contact percentage between any beads of the two residues.

### Analysis of bilayer properties

The density of phosphate beads, which was used to determine bilayer thickness, was computed using the GROMACS tool *gmx density*. To obtain the lipid lateral diffusion, jumps over periodic boundaries were removed and *gmx msd* was used upon removal of the overall center of mass motion. The second rank lipid order parameter (P_2_) was calculated for each bond of POPC using the equation

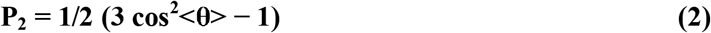

where θ represents the angle between the bilayer normal and the given bond.^35^ This was calculated using the script *do-order-gmx5*.*py* (available at www.cgmartini.nl/index.php/downloads/tools/229-do-order). The area per lipid was calculated for each bilayer leaflet separately using the membrane analysis tool Gridmat-MD, ^36^ which makes use of a grid to consider the area occupied by the TM protein. A grid with 50&50 points was used, and the area per lipid of all POPC molecules was computed every 5th frame of the trajectory.

## RESULTS

### Parameterization of DPC and characterization of DPC micelles

Due to its similarity, the parameters for DPC were mostly derived from those of POPC. Hence, choline and phosphate groups were mapped to beads of type Q1 and Q5, respectively. The hydrocarbon chain, which contains 12 carbon atoms, was described using 3 beads of type C1 (**Figure 1A**). This bead type is used in the hydrophobic tails of all saturated phospholipids in Martini 3. A force constant of 7000 kJ/mol nm^2^ was used to describe the bond between the phosphate and the choline beads, with a bond length of 0.40 nm. As this bond is present in several phospholipids, such as POPC, its force constant and length were directly obtained from the parameters for these lipids in Martini 3. For the rest of the bonds, force constants of 3800 kJ/mol nm^2^ and lengths of 0.47 nm were used. These parameters are used to describe bonds between beads in the hydrocarbon chains of all saturated phospholipids in Martini 3. The bond between the phosphate bead and the first bead of the alkyl chain was assigned the same parameters as all the bonds between beads in the tail of the lipid, as done for DPC in Martini 2. To maintain the shape of the DPC monomers, an angle of 180° between the three beads of the hydrophobic chain was maintained with a force constant of 35 kJ/mol nm^2^. Due to the similarity between the hydrocarbon chain of DPC and the hydrocarbon chains of bilayer-forming saturated phospholipids, this force constant was directly obtained from the POPC parameters.

**Figure 1:**
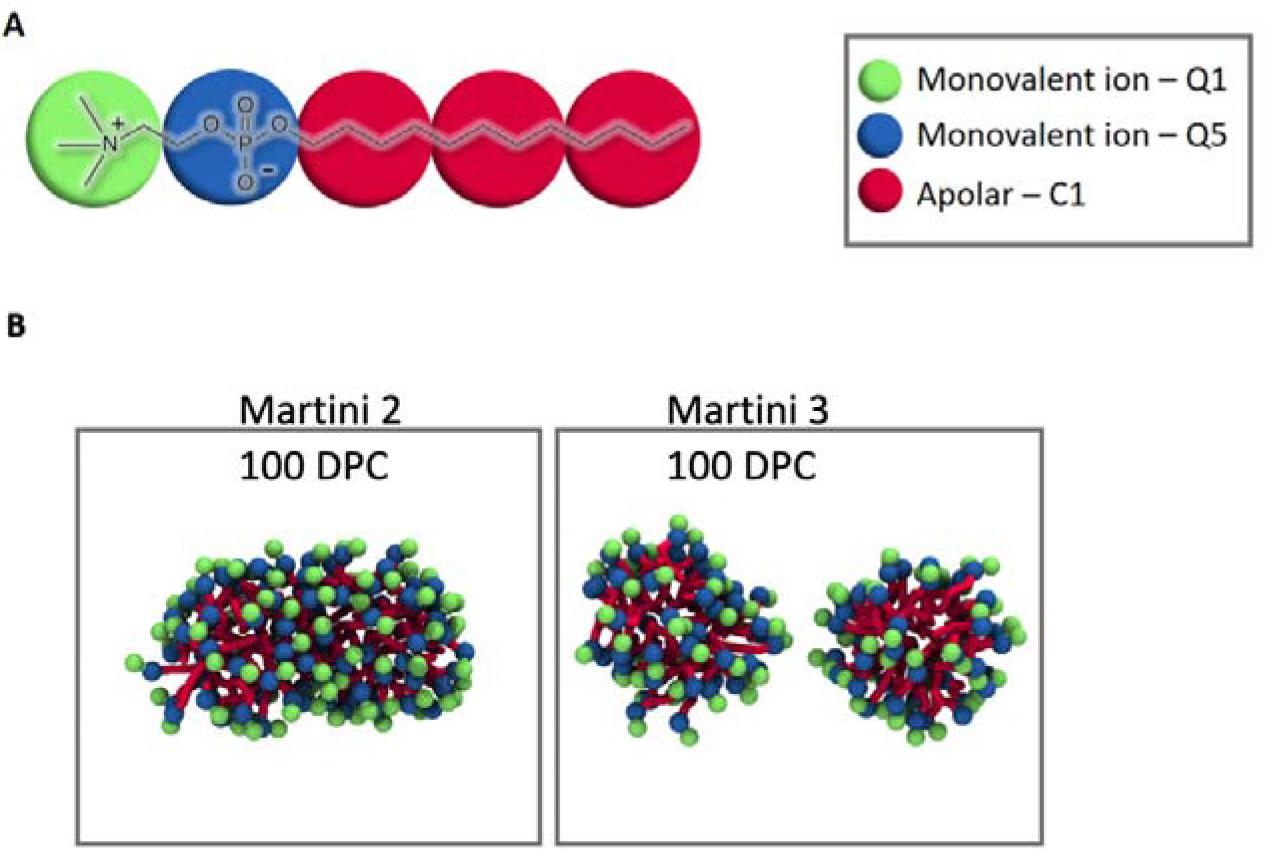
Para meterization of DPC in Martini 3. **A**: Beads of type Q1, Q5 and C1 were used to represent the choline, phosphate and hydrophobic tail groups, respectively. **B**: In simulations in a 10&10&10 nm^3^ periodic box with 100 DPC molecules (at a concentration of 166 mM) using Martini 2, a single micelle containing 100 lipids was observed. Using the same conditions and Martini 3, two smaller micelles of 45-55 molecules were obtained. **C**: Aggregation numbers obtained with Martini 3 at different DPC concentrations, compared to those obtained with Martini 2 (labelled M2) and experiments.

To investigate the ability of DPC molecules to assemble into micelles, simulations were run at different lipid concentrations (**Table 1**). Simulations using Martini 2 were computed as well in order to compare the behaviour of the preliminary DPC parameters to those of the previous version of the force field, which have been made publicly available by its developers (and were obtained from http://www.cgmartini.nl/index.php/force-field-parameters/lipids).

**Table 1:** Summary of micelle assembly simulations with different DPC concentrations. For each system, 2 replicas of 5 *µ*s each were run at 310 K using isotropic pressure coupling. The number of lipids in each aggregate at the end of each trajectory are reported for both replicas in the last column (note that these numbers vary during the simulation time).

| Martini version | # DPC | [DPC] (mM) | Box size (nm) | # Micelles | Aggregation numbers |
| --- | --- | --- | --- | --- | --- |
| Martini 3 | 100 | 166 | 10 x 10 x 10 | 2 | 43 & 57; 42 & 58 |
| Martini 3 | 120 | 199 | 10 x 10 x 10 | 2 | 44 & 76; 42 & 78 |
| Martini 3 | 140 | 233 | 10 x 10 x 10 | 2 | 60 & 80; 42 & 98 |
| Martini 3 | 160 | 266 | 10 x 10 x 10 | 3 | 40, 55 & 65; 59, 42 & 59 |
| Martini 3 | 180 | 299 | 10 x 10 x 10 | 3 | 60, 60 & 60; 62, 53 & 65 |
| Martini 3 | 200 | 333 | 10 x 10 x 10 | 3 or 4 | 45, 55, 50 & 50; 52, 50, 98 |
| Martini 3 | 100 | 125 | 11 x 11 x 11 | 2 | 42 & 58; 44 & 56 |
| Martini 3 | 100 | 96 | 12 x 12 x 12 | 2 | 43 & 57; 43 & 57 |
| Martini 3 | 100 | 76 | 13 x 13 x 13 | 2 | 43 & 57; 44 & 56 |
| Martini 2.2 | 100 | 166 | 10 x 10 x 10 | 1 | 100; 100 |
| Martini 2.2 | 100 | 100 | 12 x 12 x 12 | 1 | 100; 100 |

The most remarkable difference between DPC micelles in Martini 2 and Martini 3 was their aggregation number: whereas simulations with 100 DPC monomers (166 mM) showed the formation of a single, big micelle in Martini 2, two smaller micelles were formed in Martini 3. The distribution of lipids between them was uneven, leading to one consisting of about 57 and the other about 43 molecules (**Figure 1B**).

In simulations with 120 DPC molecules in Martini 3, formation of two micelles was observed as well; in this case, however, one micelle was significantly bigger than the other, with ca. 76 lipids. When 140 DPC molecules were included in the simulation box, two micelles with around 60 and 80 lipids were formed. 160 DPC monomers assembled into 3 micelles with ~ 40, 55 and 65 molecules, and 180 lipids formed 3 more uniform micelles with ~ 60 monomers each. When the DPC concentration was further increased up to 333 mM (200 DPC molecules), 4 micelles were formed. The polar groups of adjacent micelles tended to interact with each other, eventually leading to the fusion of two micelles into elongated, tubular structures with ~ 98 monomers each.

In general, the aggregation numbers obtained with the derived DPC parameters in Martini 3 show better agreement with experimental values than those obtained using Martini 2. Although the DPC aggregation numbers described in the literature vary considerably, which can be due to differences in the method, temperature or lipid concentration, they generally agree with the results obtained here. NMR spectroscopy experiments with DPC concentrations of 228 mM and 98 mM showed aggregation numbers of 44 ± 5 and 56, respectively.^12 37^ Higher aggregation numbers have been observed with small-angle X-ray scattering (SAXS, 60-70 molecules) and small-angle neutron scattering (SANS, 70.6 ± 5 molecules).^38 39^ Ultracentrifugation experiments showed aggregation numbers of 56 ± 5 at 20 mM and 294 K.^40^

Similar aggregation numbers (~ 54 molecules) have been described in atomistic MD simulations using different force fields.^41 42 43^ CG MD simulations using Martini 2 showed a high variability of aggregation numbers: at a temperature of 300 K and a concentration of 40 mM, micelles of 40 to 70 molecules have been reported,^35^ whereas at 300 K and a lipid concentration of 126 mM (175 molecules), micelles consisting of 45 monomers were formed.^44^ A summary of all DPC aggregation numbers obtained in previous experimental and simulation studies can be found in **Table 2**.

**Table 2:**
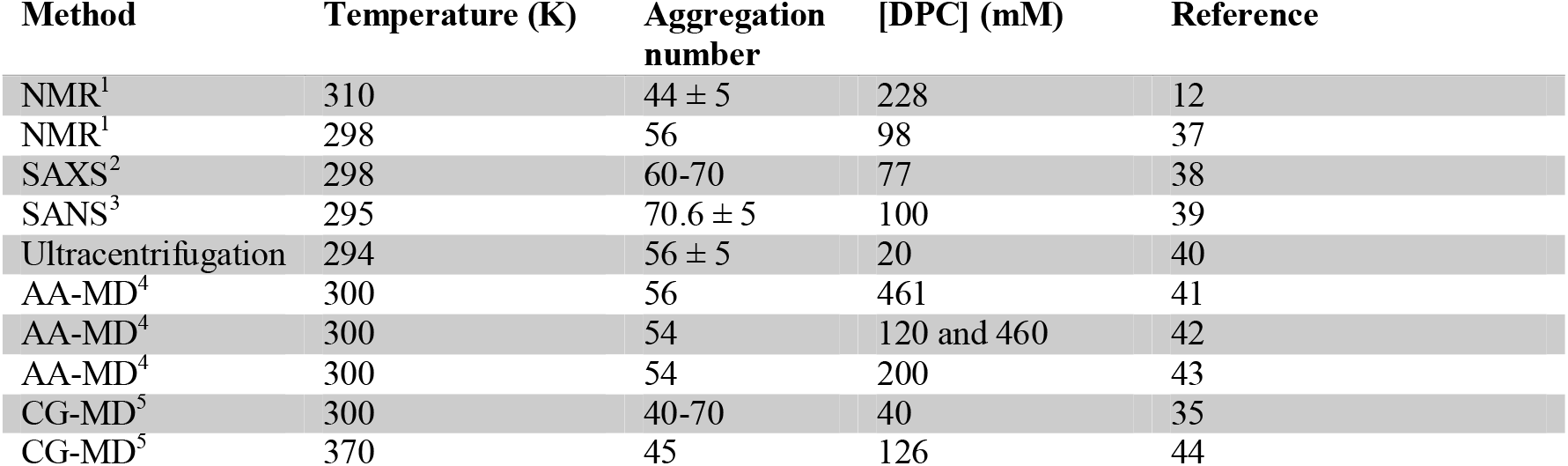
DPC aggregation numbers reported in the literature.

| Method | Temperature (K) | Aggregation number | [DPC] (mM) | Reference |
| --- | --- | --- | --- | --- |
| NMR<sup>1</sup> | 310 | $44 \pm 5$ | 228 | 12 |
| NMR<sup>1</sup> | 298 | 56 | 98 | 37 |
| SAXS<sup>2</sup> | 298 | 60-70 | 77 | 38 |
| SANS<sup>3</sup> | 295 | $70.6 \pm 5$ | 100 | 39 |
| Ultracentrifugation | 294 | $56 \pm 5$ | 20 | 40 |
| AA-MD<sup>4</sup> | 300 | 56 | 461 | 41 |
| AA-MD<sup>4</sup> | 300 | 54 | 120 and 460 | 42 |
| AA-MD<sup>4</sup> | 300 | 54 | 200 | 43 |
| CG-MD<sup>5</sup> | 300 | 40-70 | 40 | 35 |
| CG-MD<sup>5</sup> | 370 | 45 | 126 | 44 |

### Systems of transmembrane helix dimer proteins embedded in DPC micelles or in self-assembled POPC bilayers are not achievable in Martini 3

TM helix dimers are components of many receptors and are thought to play an important role in transmitting extracellular signals to the cellular interior.^45^ Experimental determination of the structure of TM helix dimers is often performed for TM helix dimers immersed in DPC micelles rather than a membrane bilayer. To investigate the insertion of TM helix dimers into

DPC micelles using the previously derived DPC parameters, simulations starting with 100 DPC molecules at random positions and a position-restrained glycophorin A (GpA) TM helix dimer were run with Martini 3. This protein is considered to represent an archetypal TM helix dimer and is often used to assess computational methods for studying TM helix dimer proteins. ^22 33 46 47^ Surprisingly, although the DPC lipids assembled into micelles successfully, with aggregation numbers similar to those obtained in simulations without protein, GpA was not encapsulated within a micelle. The TM helix dimer interacted with the polar head beads of the DPC lipids and with the water beads rather than with the hydrocarbon chains (**Figure 2A**).

**Figure 2:**
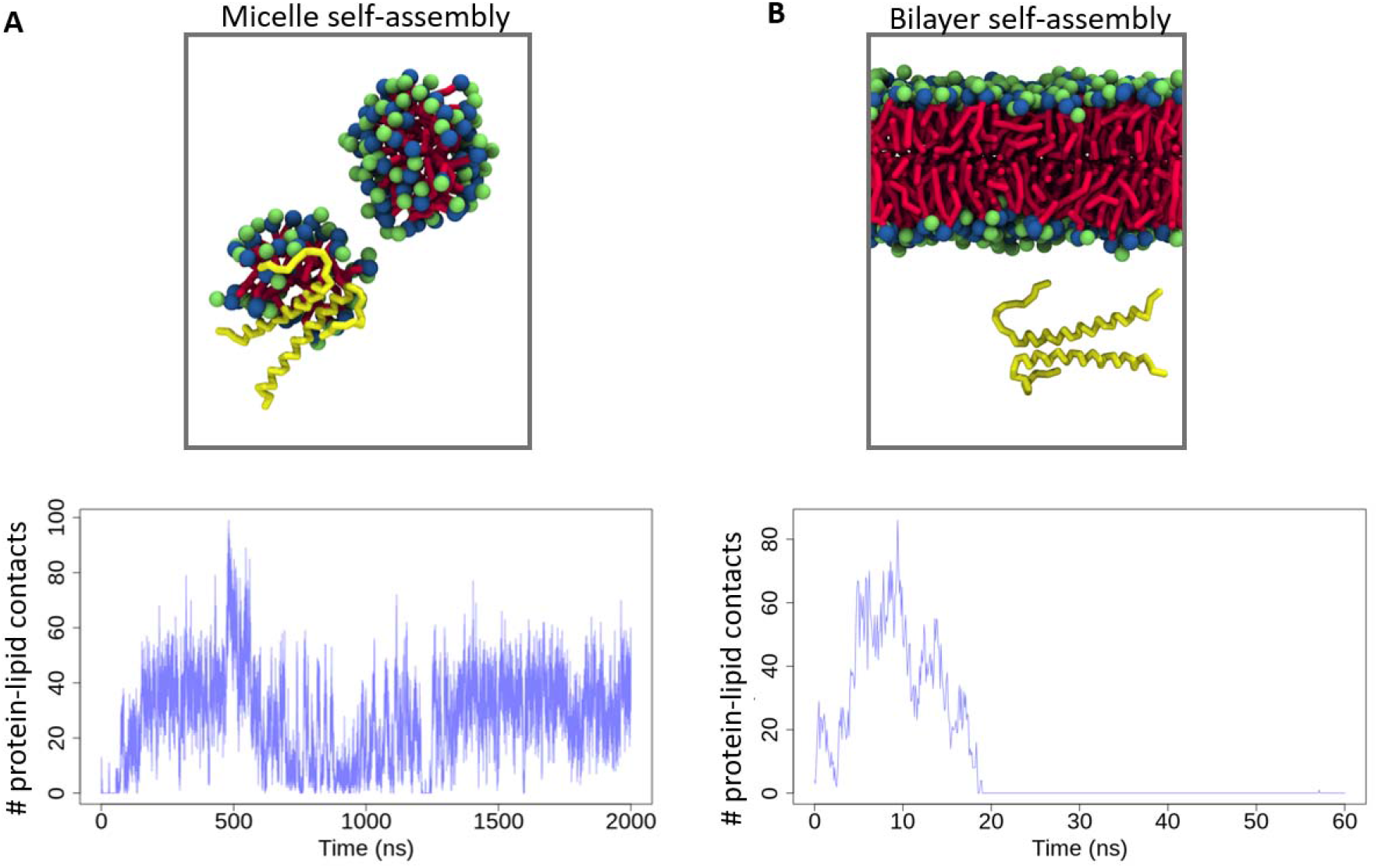
Aberrant behaviour of transmembrane helix dimer proteins in lipid environments simulated using Martini 3. **A**: GpA TM helix dimers (yellow) do not interact with the hydrophobic tails (red) of micelle-forming DPC molecules, but rather with the water and with the polar head groups (blue and green) of the lipids. **B**, The POPC bilayer does not assemble around the position-restrained GpA, resulting in the absence of contacts between the protein and the lipid.

To eliminate a possible sequence dependence of this effect, simulations in identical conditions were run for the TM helix dimers of the neurotrophin receptor TrkA. Similarly, a lack of insertion in micelles was observed. In simulations starting from a micelle-embedded TM helix dimer, immediate micelle dissociation and re-assembly excluding the protein from the micelles was observed (**Supplementary Figure 1**).

We considered that a wrong choice of DPC parameters could be the reason for the lack of protein-lipid interactions in these simulations, given that these parameters have not been as extensively validated as those for other phospholipids that have been released by the Martini 3 developers. To test this hypothesis, simulations of the GpA dimer with and without position restraints and POPC lipids at random initial positions were run. The rationale behind these tests is that this lipid has been parameterized and validated by the developers of the Martini 3 force field; thus, protein-lipid interactions should be correctly reproduced in systems containing POPC. Strikingly, although the lipid molecules rapidly assembled to form a bilayer, GpA was not embedded inside it but rather interacted with the water beads (**Figure 2B**). In those simulations in which the protein dimer was not position-restrained during bilayer self-assembly, the two monomers showed a tendency to dissociate from one another (**Supplementary Figure 2**). On the other hand, in simulations that had been prepared using the INSANE method to insert transmembrane proteins into preassembled bilayers, the GpA TM helix dimers remained inside the membrane for the entire trajectory (20 μs).

### Scaling down protein-water interactions in Martini 3 enables simulation of transmembrane helix dimer proteins in DPC micelles and in self-assembled phospholipid bilayers

Different strategies were tested in order to correct the lack of interaction between TM helix dimers and the hydrophobic tails of DPC and POPC molecules. The first group of strategies did not use any modification of the interaction parameters of the Martini 3 force field. These approaches mainly focused on the insertion of the transmembrane proteins into micelles, and were based on modifications of the previously described DPC parameters. A more detailed description of these tests, which were conducted using TrkA TM helix dimers, can be found in **Supplementary Table 1 and Supplementary Figure 3**. None of these strategies led to insertion of the protein into micelles.

The second group of strategies was based on the modification of the nonbonded parameters for interactions between given bead types (as described in the Methods section). In order to correct for the lack of protein-lipid interactions, non-bonded interactions between protein beads and lipid chains were scaled up as described in the Methods section, using a values greater than one. Although a slight increase in protein-lipid interactions was observed, this was not sufficient to fully embed the protein into the micelles. Furthermore, after release of the position restraints, the protein monomers dissociated from one another in all replicas and either ended in different micelles or on opposite sides of the same micelle (**Supplementary Figure 3**).

In view of the lack of success of an increase of protein-lipid non-bonded interactions, a different strategy was tested: decreasing protein-water interactions. Applying a scaling factor a of 0.9 to all interactions between protein and water beads, 10 replicas of 20 *µ*s each were run for a system containing 100 DPC molecules and a GpA TM helix dimer. In all replicas, a large micelle with 100 lipids assembled around the protein (**Figure 3A**) and remained stable during the entire trajectory. Similar results were observed when a TrkA TM helix dimer was simulated under the same conditions.

**Figure 3:**
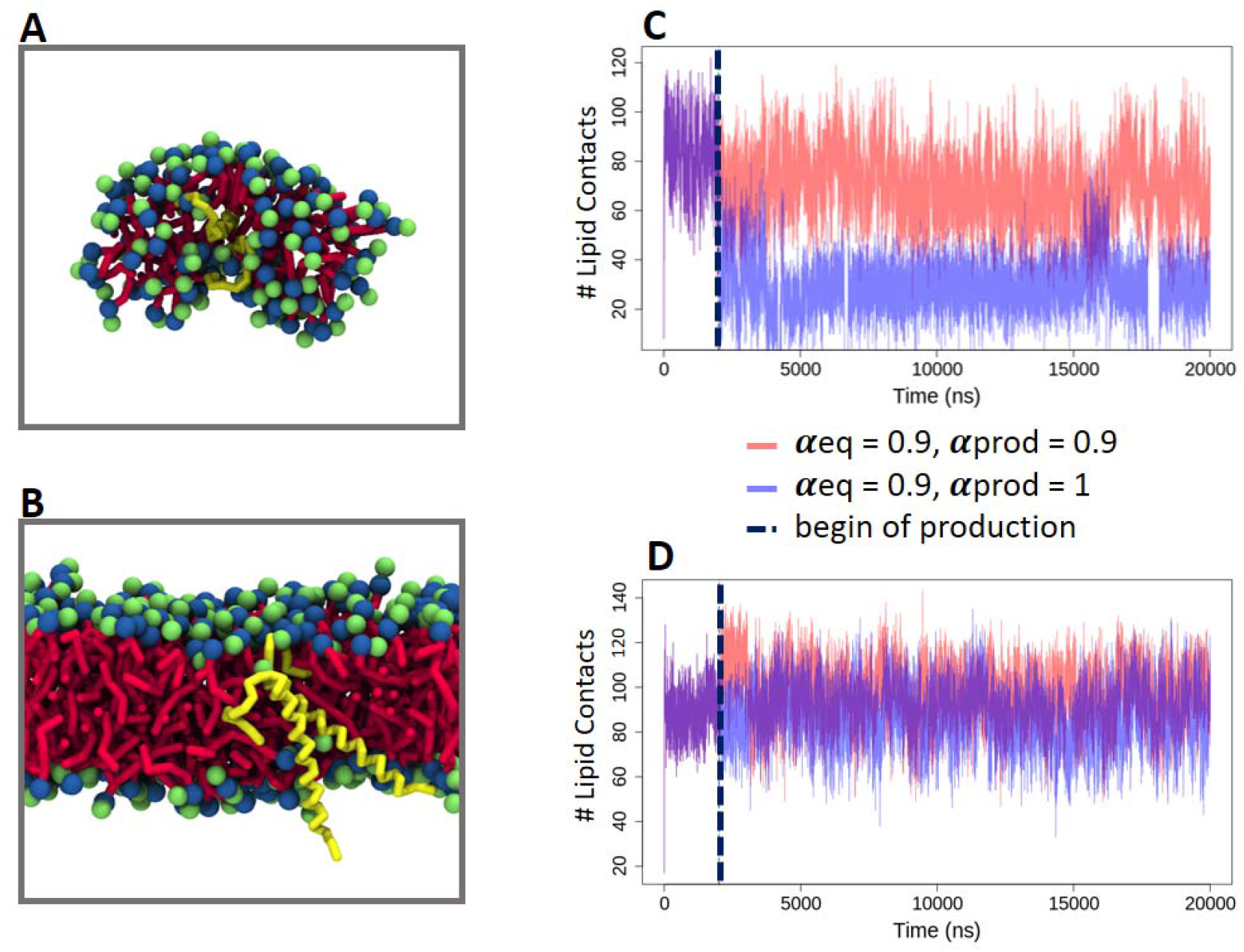
Protein-lipid systems simulated with Martini 3 after introduction of a scaling factor. **A, B**: Decreasing protein-water interactions by 10% (α = 0.9) leads to correct insertion of GpA dimers into a micelle (A) and a POPC bilayer (B). **C**: After removal of the scaling factor at the beginning of the production simulation (dashed line), GpA leaves the micelle, which results in a reduction in the number of protein-lipid contacts (blue line) compared to maintaining the scaling factor (red line). **D**: In simulations of GpA dimers in POPC bilayers, the protein dimer remains inside the membrane upon removal of the scaling factor after equilibration.

Removing the scaling after an equilibration of 2 *µ*s led to immediate dissociation of the large micelle containing 100 DPC molecules into two smaller ones and interruption of the interactions with the transmembrane protein (**Figure 3C**). This suggests that the scaling factor proposed here needs to be maintained during the totality of the trajectory in order to simulate TM proteins in micelles.

Scaling down protein-water interactions by 10% also led to POPC bilayer self-assembly around TM helix dimers (**Figure 3B**). In order to assess the statistical robustness of these observations, 100 short (60 ns) simulations of bilayer self-assembly in the presence of GpA were performed with different scaling factors. Whereas assembly of the phospholipid bilayer around the TM helix dimer was observed in only one out of 100 simulations without scaling, it was observed in 30 trajectories when a scaling factor of *α* = 0.9 was used (**Supplementary Figure 4**).

10 of the 100 short simulations with *α* = 0.9 were equilibrated for 2 *µ*s. Afterwards, each of them was used as input for two 20 *µ*s production runs: one maintaining protein-water scaling, and the other having the default Martini 3 parameters. In half of the replicas, the protein had acquired a correct orientation inside the lipid bilayer during equilibration. In all these cases, the protein remained inside the membrane during the production run regardless of whether it was run with or without down-scaling of the protein-water interactions (data from two trajectories are shown in **Figure 3D**). Thus, in contrast to simulations of TM helix dimers in micelles, where scaling seems to be crucial to obtain meaningful results, in the case of self-assembled bilayers, scaling can be removed as soon as the TM helix dimer acquires the correct orientation in the bilayer.

In the remaining half of the replicas, GpA did not orient correctly inside the membrane, acquiring either a diagonal or a horizontal arrangement, with one of the two monomers partially outside of the bilayer. In these cases, the protein exited the membrane after release of the position restraints in production runs. This behaviour was observed in simulations with and without scaling of protein-water interactions.

To assess the general applicability of scaling the protein-water interactions during self-assembly, we ran a further set of simulations of membrane self-assembly in the presence of a GpA TM dimer using DPPC instead of POPC. 10 replica simulations, each 2 *µ*s long, were first run using the original Martini 3 parameters and then run with the protein-water interactions scaled down by 10%. In accordance with the preceding results, protein insertion in the bilayer was only observed in one out of the ten replicas using the unscaled Martini 3 parameters, while scaling the protein-water interactions led to GpA being inserted in the DPPC bilayer in 8 replicas (**Supplementary Figure 5**).

### Scaling of protein-water interactions does not affect the dimerization behaviour of transmembrane helix dimers

To investigate whether the down-scaling of protein-water interactions affects the dimer arrangements adopted by GpA in a membrane, structural and contact information was extracted from the simulations of the GpA dimer in self-assembled POPC bilayers with and without scaling described in the previous section. Only those replicas (5 out of 10) in which the TM helix dimer remained inside the lipid bilayer during the entire trajectory were used for this analysis.

To analyse the conformational landscape of the GpA dimer, all five replicas were merged and the whole system was aligned to one of the two helices. A series of structural parameters was computed to characterize the different possible helix dimer arrangements. These parameters included the distance between the centres of geometry (COG), the crossing angle, and the rotation angles around the first and second helices (**Figure 4A**, also described in the Methods section). Kernel density estimation (KDE) was used to identify the most populated conformations as a two-dimensional function of the given structural parameters.

**Figure 4:**
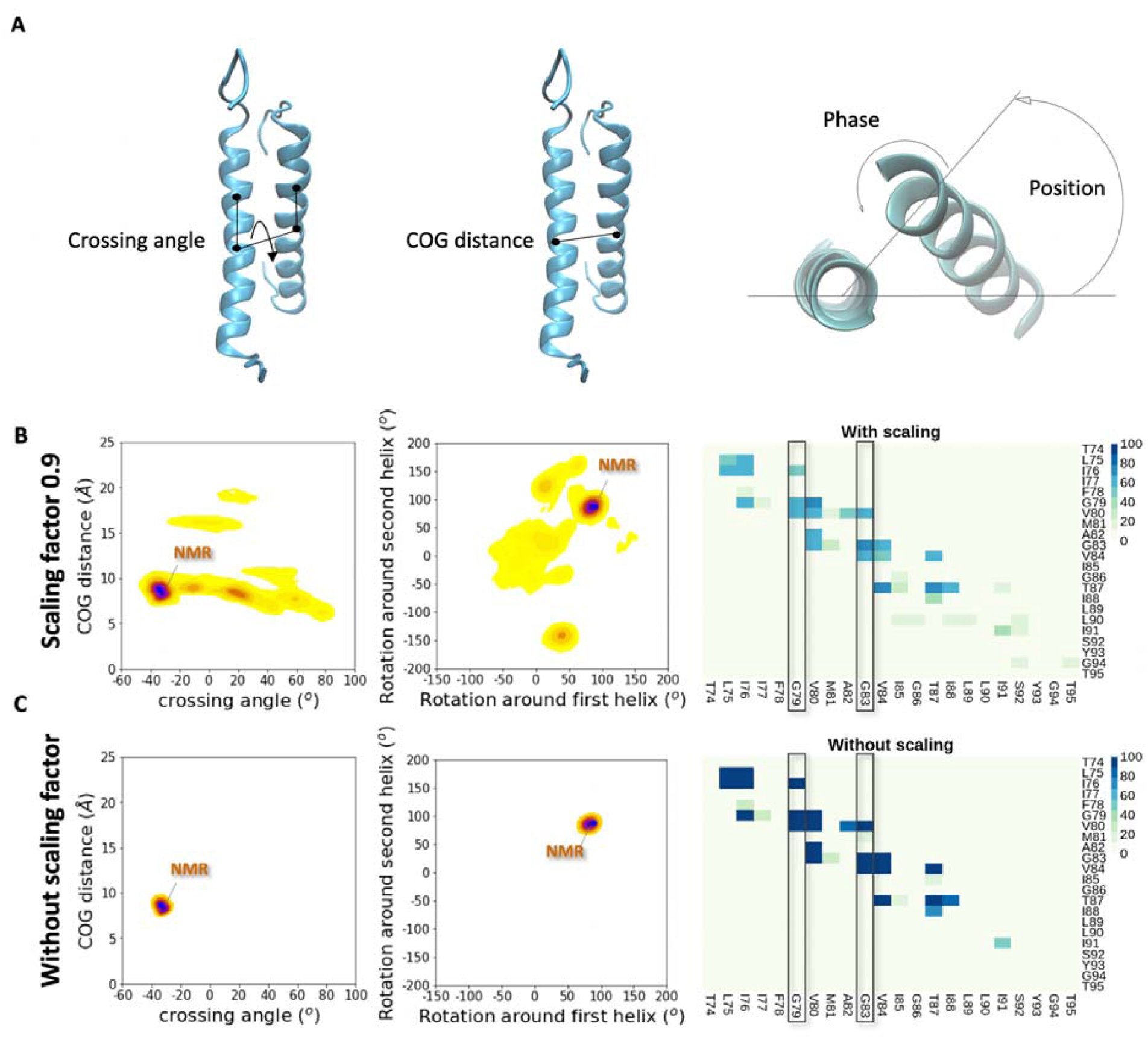
Transmembrane helix dimer proteins conserve their structural properties upon scaling protein-water interactions down by 10% in Martini 3. **A**: Structural parameters used to define the conformational space of helical dimers, including the crossing angle, distance between centers of geometry (COG) of both helices, position and phase angles. Position and phase angles represent the rotations of the non-centered (“second”) helix with respect to the centered (“first”) helix and with respect to itself, respectively (see **Methods**). **B**: Conformational space and contact analysis of GpA TM helix dimers in POPC bilayers simulated with a uniform scaling factor of = 0.9. Kernel density estimation was used to determine the most populated conformations described by the crossing angle and the COG distance (left panel) or the rotation dihedrals around the two helices (middle panel). This analysis was run on 5 merged replicas, each 20 μs long. Positive and negative crossing angles denote left- and right-handed dimers, respectively. The blue dot corresponds to the NMR structure, which is at a density maximum in both representations of the conformational landscape. The dimerization motif ^79^*GxxxG*^83^ is part of the dimer interface during most of the simulation time (right panel). A cut-off distance of 6 Å was used to define contacts and their percentage occupation is given by color in the heat map. **B**: Conformational space and contact analysis of GpA TM helix dimers in POPC bilayers simulated without introducing a scaling factor. The explored portion of the conformational space is narrower than in simulations with a scaling factor, with a single density maximum corresponding to the NMR structure. The dimer interface contains the same residues as in simulations with a scaling factor, including the ^79^*GxxxG*^83^ dimerization motif.

A greater exploration of the conformational landscape was observed in simulations in which protein-water interactions had been reduced by 10% (**Figure 4B, Supplementary Figure 6**). However, the density maxima corresponding to the conformations of the NMR structure of GpA were observed in simulations both with and without scaling of protein-water interactions (**Figure 4C**). These helical arrangements showed a crossing angle of about -34º, a distance between COGs of 8-9 Å, and position and phase angles around 90º with Gly 79 defined as the reference residue. A similar conformational landscape was observed in simulations of GpA dimers inserted into a preformed membrane with the INSANE method without protein-water scaling (**Supplementary Figure 7**). Hence, down-scaling protein-water interactions does not seem to affect the structural arrangements of the GpA TM helix dimers.

To further investigate the effect of the scaling of protein-water interactions on the dimerization behaviour of GpA, contact analysis was performed for simulations performed with and without the introduction of the scaling factor. In both cases, the same residues participated in the dimer interface, which contained the dimerization motif ^79^GxxxG^83^. This sequence has been proposed to be crucial for the dimerization of GpA. Residues L75, I76, V80, G83, V84 and T87 were part of the dimer interface as well during most of the trajectories (**Figure 4B and 4C**). Thus, it can be concluded that introducing a factor to down-scale protein-water interactions by 10% does not disturb the dimer interface or other structural parameters of TM helix dimers embedded in lipid bilayers.

To evaluate the effect of the scaling on the POPC bilayers, structural parameters of the lipid bilayer were calculated for simulations with and without scaling. These include the lipid lateral diffusion, phosphate density (which gives an estimate of the bilayer thickness), area per lipid, and second rank order parameter (P_2_). No noticeable differences in these parameters were observed between simulations performed with and without scaling (**Supplementary Figure 8**), which indicates that the reduction of the protein-water interactions does not affect the structure of the phospholipid bilayer.

The conformational landscape of the GpA TM helix dimers in DPC micelles in simulations with a 10% reduction of protein-water interactions showed notable differences to that observed in POPC bilayers (**Figure 5**). This indicates a significant effect of the lipid environment on the structure of TM helix dimers in Martini 3.

**Figure 5:**
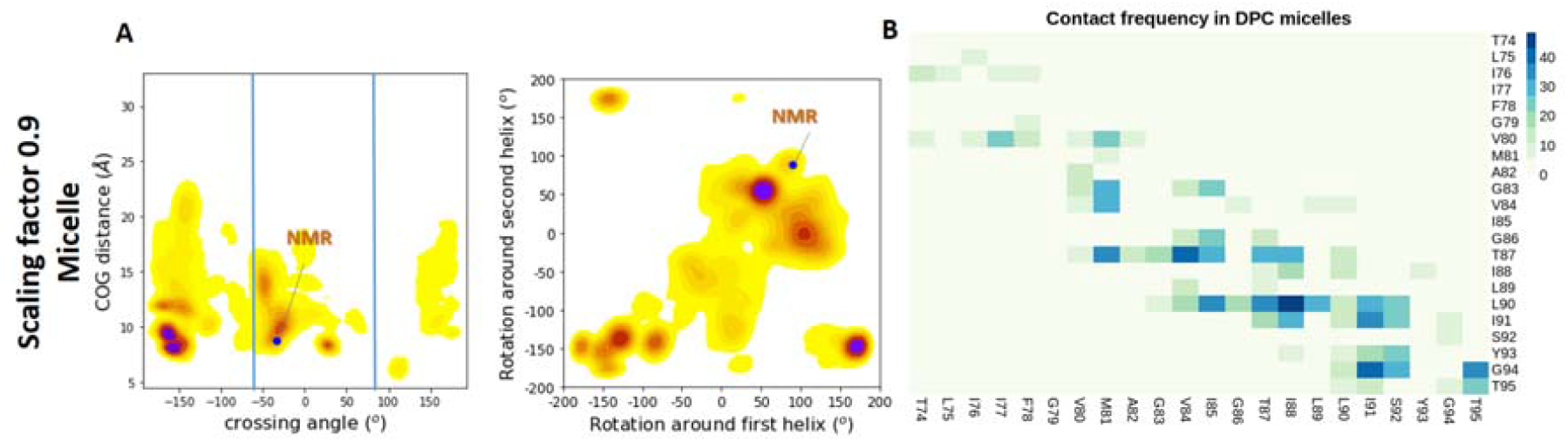
Structural features of GpA dimers in DPC micelles upon reduction of protein-water interactions by 10% in Martini 3. **A**: A significantly greater conformational space is explored compared to simulations in POPC bilayers (shown in figure 4). In many frames, the dimer adopts higher or lower crossing angles than observed in bilayers (beyond the blue vertical lines). This analysis was run on 10 merged replicas, each 20 μs long. The blue dot represents the NMR structure. **B**: Percentage occupancy of residue-residue contacts between the two TM helix monomers in DPC micelles. Dimers with crossing angles greater than 80º and less than -60º were not considered for this analysis due to their lack of relevance in biological membranes. A cut-off distance of 6 Å was used to define contacts.

## DISCUSSION

Detergent micelles are commonly used to study the structural properties of TM proteins. MD simulations are usually carried out for such proteins in phospholipid bilayers, which better resemble their physiological environment. However, this means that it is not always clear whether differences between simulations and experiments arise from the different nature of these methods or from the differences in the lipid environments. Simulation of TM proteins in detergent micelles can thus help to clarify this matter, and if combined with simulations in bilayers, enable comparison of experimental and computational results.

In the present study, we show that TM helix dimers cannot be simulated in DPC micelles using the most recent version of the Martini force field, Martini 3, due to a lack of interaction of the protein with the hydrophobic tails of the surfactant molecules. We also demonstrate that self-assembled phospholipid bilayers rarely encapsulate TM helix proteins in Martini 3. These observations suggest that the TM protein beads are relatively hydrophilic in Martini 3. Indeed, a comparison of the Lennard Jones potentials of beads of different residues with hydrophobic lipid tail beads and with water beads shows remarkable differences between hydrophilic and hydrophobic amino acids (**Supplementary Figure 9**).

When TM helix dimers are inserted in preformed membranes using the INSANE method, however, they behave as expected and remain inside the lipid bilayer. This is consistent with previously reported studies of other TM helix dimers using Martini 3, e.g. of Ephrin type-A receptor I (EphA1) dimerization.^48 49^ On the contrary, when simulations were initiated with a TM helix dimer inserted in a preformed DPC micelle, the micelle dissociated immediately, leaving the protein exposed to water. This difference between bilayer and micelle environments can arise because the former is more stable, while the latter is more dynamic and thus the protein has more degrees of freedom.^50 51^

Here, we show that introducing a scaling factor to reduce protein-water interactions by 10% in Martini 3 results in insertion of GpA TM helix dimers into phospholipid bilayers and detergent micelles. The fact that Martini 3 uses a specific bead type for water, which has been parameterised independently, makes this solution very attractive. Indeed, we find that this down-scaling does not affect the dimerization behaviour of the GpA TM helices. The structural and dynamic parameters of the lipid bilayers, such as bilayer thickness and lateral diffusion, also remain unaltered upon scaling.

Protein-water interactions have recently been scaled up in Martini 3 to reproduce the experimental dimensions of soluble and intrinsically disordered proteins.^11^ The fact that opposite adjustments need to be made to reproduce the experimental behaviour of TM helix dimers reflects the challenging nature of CG force field parameterization: due to a reduction of the chemical space, system-dependent adjustments need to be made in order to obtain biologically meaningful results. Indeed, scaling of the Lennard-Jones parameters has been widely used to adapt the previous version of the force field to different biomolecular systems.^52 53^

We observe remarkable differences in the conformational landscapes of GpA dimers in POPC bilayers and DPC micelles. We believe these differences are due to the more restraining environment of POPC bilayers as opposed to the more dynamic nature of micelles, which allow the helix dimers to adopt arrangements with much larger crossing angles. Indeed, it has been shown experimentally that the fluctuations in the crossing angles of GpA dimers in DPC micelles are twice as large as those in bilayers.^54^ Another study suggests that the stability of GpA dimers increases linearly with the aggregation number of the detergent.^55^ As the aggregation number of DPC micelles is significantly lower with our Martini 3 parameters than with Martini 2, one would expect the stability of the TM helix dimer to be lower as well. Other peptides have been shown to adopt differing conformations in DPC micelles and disk-like bicelles. ^51 56 57^ We believe the differences in conformational landscape in micelles and bilayers found in this study are caused by the differences in lipid environment rather than by the scaling applied to the protein-water interactions. The fact that only minimal differences are observed in the helix dimer arrangements obtained in bilayers with and without scaling supports this hypothesis. Hence, we expect the scaling of the non-bonded protein-water interactions in Martini 3 proposed here to provide a rapid, simple and effective correction of the aberrant behaviour of TM helix proteins in self-assembled lipid bilayers and detergent micelles.

We recommend using this adjustment to the Martini 3 force field in all simulations of transmembrane proteins that rely on membrane or micelle self-assembly. For membrane systems, once the protein has been encapsulated in the membrane, the scaled parameters can be replaced by the original ones, as we did not observe aberrant behaviour of the TM proteins once they were inserted in the lipid bilayer. Nevertheless, simulations of TM proteins using the original Martini 3 parameters should be carefully examined. Furthermore, we advise users of Martini 3 to apply this scaling in simulations of TM proteins in micelles, as this type of system cannot be simulated using the original Martini 3 force field.

Although changing the protein−water interaction strength may affect other properties such as oil/water transfer energies, the changes we propose are still small relative to the large free energies typically associated with bilayer insertion.

Moreover, we have also shown that important properties of transmembrane protein systems, such as the structural parameters of bilayers and protein dimers, are well preserved. Hence, if this scaling is applied in the context we propose, one can obtain biologically meaningful results without compromising the general behavior of the optimised Martini 3 force field parameters.

Lastly, we expect these results to be useful for the further optimization of the Martini 3 force field. In particular, our data suggest that tests in micelles should be included in the parameterisation procedure, as they provide a sensitive test of the correct behavior of protein– lipid and protein–water interactions.

## ACKNOWLEDGEMENTS

This research was funded by the European Union’s Horizon 2020 research and innovation programme “Euroneurotrophin” under the Marie Skłodowska-Curie grant agreement No 765704 and the Klaus Tschira Foundation. The authors acknowledge support by the state of Baden-Württemberg through bwHPC and the German Research Foundation (DFG) through grant INST 35/1134-1 FUGG. We thank Paulo Cesar Telles de Souza for helpful discussions and Stefan Richter for technical support.

## DATA AVAILABILITY

The trajectories generated for this study are freely available in Zenodo at the DOI: 10.5281/zenodo.7061501.

## ASSOCIATED CONTENT

Supporting Information available: Supplementary Table 1, summary of all simulations run in this study; Supplementary Figure 1, lack of interactions between TM TrkA dimers and the hydrophobic tails of DPC molecules in simulations with the standard Martini 3 parameters; Supplementary Figure 2, GpA monomers that are not embedded into the lipid bilayer tend to dissociate; Supplementary Figure 3, different unsuccessful approaches to increase protein-lipid interactions and achieve insertion of the TrkA dimer into micelles; Supplementary Figure 4, number of contacts between GpA and POPC in simulations with different values of the scaling factor for protein-water non-bonded interactions; Supplementary Figure 5, number of contacts between GpA and the hydrophobic tails of the DPPC lipids during self-assembly simulations with and without scaling of the protein-water interactions in Martini 3; Supplementary Figure 6, root mean square deviation (RMSD) of the GpA dimer in different replicas with and without application of the scaling factor of α = 0.9 to reduce protein-water non-bonded interactions; Supplementary Figure 7, Conformational landscape of GpA TM helix dimers in POPC bilayers inserted in the membrane via the *INSANE* method and simulated without introduction of a scaling factor using Martini 3; Supplementary Figure 8, structural properties of the POPC phospholipid bilayers do not differ in simulations using Martini 3 with and without application of the scaling factor of α = 0.9 to reduce protein-water non-bonded interactions; Supplementary Figure 9, Lennard-Jones potentials of beads of selected residues with hydrophobic and water beads in Martini 3.

Supplementary Files: DPC structure and parameters, Gromacs mdp parameter files for different systems, Martini 3 force field parameters after scaling down protein-water interactions, R script which was used to scale down protein-water interactions, TCL script to find interhelical contacts during a given trajectory, R scripts which were used to perform different analysis tasks, contact data.

## Footnotes

1 Nuclear Magnetic Resonance

2 Small-Angle X-ray Scattering

3 Small-Angle Neutron Scattering

4 All-Atom Molecular Dynamics simulation

5 Coarse-Grained Molecular Dynamics

