## Supplementary Figures and Tables for "Scaling protein-water interactions in the Martini 3 coarse-grained force field to simulate transmembrane helix dimers in different lipid environments"

### Supplementary information

*Ainara Claveras Cabezudo*[†,‡], *Christina Athanasiou*[†,‡,§], *Alexandros Tsengenes*[†,‡,§], *Rebecca C. Wade*[*,†,‡,‖,⊥]

† Molecular and Cellular Modeling Group, Heidelberg Institute for Theoretical Studies (HITS), Schloss-Wolfsbrunnenweg 35, 69118 Heidelberg, Germany

‡ Faculty of Biosciences, Heidelberg University, Im Neuenheimer Feld 234, 69120 Heidelberg, Germany

§ Heidelberg Biosciences International Graduate School, Heidelberg University, Im Neuenheimer Feld 501, 69120 Heidelberg, Germany

‖ Center for Molecular Biology (ZMBH), DKFZ-ZMBH Alliance, Heidelberg University, Im Neuenheimer Feld 282, 69120 Heidelberg, Germany

⊥ Interdisciplinary Center for Scientific Computing (IWR), Heidelberg University, Im Neuenheimer Feld 205, 69120 Heidelberg, Germany

*

### Supplementary Table 1: Summary of all simulations run in this study.

| Lipid system | Protein | Martini version | No. of Replicas | Duration of each replica (μs) | No. of Lipids | Lipid concentration (mM) | Box size (nm) |
|---|---|---|---|---|---|---|---|
| DPC (self-assembly) | - | Martini 3 | 2 | 5 | 100 | 166 | 10 x 10 x 10 |
| DPC (self-assembly) | - | Martini 3 | 2 | 5 | 120 | 199 | 10 x 10 x 10 |
| DPC (self-assembly) | - | Martini 3 | 2 | 5 | 140 | 233 | 10 x 10 x 10 |
| DPC (self-assembly) | - | Martini 3 | 2 | 5 | 160 | 266 | 10 x 10 x 10 |
| DPC (self-assembly) | - | Martini 3 | 2 | 5 | 180 | 299 | 10 x 10 x 10 |
| DPC (self-assembly) | - | Martini 3 | 2 | 5 | 200 | 333 | 10 x 10 x 10 |
| DPC (self-assembly) | - | Martini 3 | 2 | 5 | 100 | 125 | 11 x 11 x 11 |
| DPC (self-assembly) | - | Martini 3 | 2 | 5 | 100 | 96 | 12 x 12 x 12 |
| DPC (self-assembly) | - | Martini 3 | 2 | 5 | 100 | 76 | 13 x 13 x 13 |
| DPC (self-assembly) | - | Martini 2.2 | 2 | 5 | 100 | 166 | 10 x 10 x 10 |
| DPC (self-assembly) | - | Martini 2.2 | 2 | 5 | 100 | 100 | 12 x 12 x 12 |
| DPC (self-assembly) | TrkA | Martini 3 | 4 | 7 | 100 | 166 | 10 x 10 x 10 |
| DPC (start with protein inserted in micelle) | TrkA | Martini 3 | 2 | 5 | 100 | 166 | 10 x 10 x 10 |
| DPC-C1r (self-assembly) | TrkA | Martini 3 | 5 | 5 | 100 | 166 | 10 x 10 x 10 |
| DPC-SC1 (self-assembly) | TrkA | Martini 3 | 5 | 5 | 100 | 166 | 10 x 10 x 10 |
| DPC (self-assembly) | TrkA | Martini 3 (α for protein-lipid = 1.08) | 5 | 20 | 100 | 166 | 10 x 10 x 10 |
| DPC (self-assembly) | TrkA | Martini 3 (α for protein-lipid = 1.04) | 5 | 20 | 100 | 166 | 10 x 10 x 10 |
| DPC (self-assembly) | TrkA | Martini 3 (α for protein-lipid = 1.09) | 5 | 20 | 100 | 166 | 10 x 10 x 10 |

| | | | | | | | |
|---|---|---|---|---|---|---|---|
| DPC (self-assembly) | TrkA | Martini 3 (α for protein-lipid = 1.09) | 5 | 20 | 100 | 166 | 10 x 10 x 10 |
| DPC (self-assembly) | GpA | Martini 3 | 10 | 20 | 100 | 166 | 10 x 10 x 10 |
| DPC (self-assembly) | GpA | Martini 3 (α for protein-lipid = 1.08) | 5 s | 20 | 100 | 166 | 10 x 10 x 10 |
| DPC (self-assembly) | GpA | Martini 3 (α for protein-water = 0.9) | 10 | 20 | 100 | 166 | 10 x 10 x 10 |
| POPC (self-assembly) | GpA | Martini 3 | 10 | 20 | 250 | 415 | 10 x 10 x 10 |
| POPC (self-assembly) | GpA | Martini 3 | 100 | 0.06 | 250 | 415 | 10 x 10 x 10 |
| POPC (self-assembly) | GpA | Martini 3 (α for protein-water = 0.9) | 100 | 0.06 | 250 | 415 | 10 x 10 x 10 |
| POPC (self-assembly) | GpA | Martini 3 (α for protein-water = 0.93) | 100 | 0.06 | 250 | 415 | 10 x 10 x 10 |
| POPC (self-assembly) | GpA | Martini 3 (α for protein-water = 0.95) | 100 | 0.06 | 250 | 415 | 10 x 10 x 10 |
| POPC (self-assembly) | GpA | Martini 3 (α for protein-water = 0.9) | 10 | 20 | 250 | 415 | 10 x 10 x 10 |
| POPC (self-assembly) | GpA | Martini 3 | 10 | 20 | 250 | 415 | 10 x 10 x 10 |
| DPPC (self-assembly) | GpA | Martini 3 (α for protein-water = 0.9) | 10 | 2 | 250 | 415 | 10 x 10 x 10 |
| DPPC (self-assembly) | GpA | Martini 3 | 10 | 2 | 250 | 415 | 10 x 10 x 10 |
| POPC (*INSANE*) | GpA | Martini 3 | 10 | 20 | 185 | 436 | 8 x 8 x 11 |

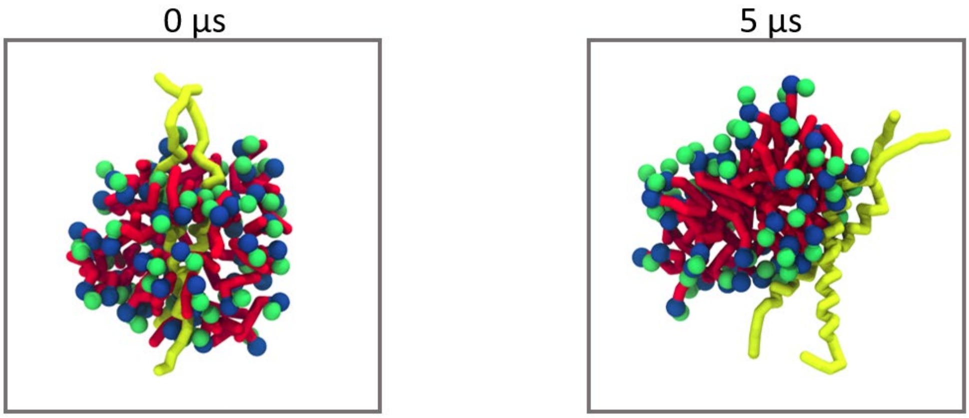


**Supplementary Figure 1: Lack of interactions between TM TrkA dimers and the hydrophobic tails of DPC molecules in simulations with the standard Martini 3 parameters.** Simulations beginning with a position-restrained TM helix dimer embedded in a DPC micelle, as shown on the left, were run. During simulation, the DPC micelle encapsulating the TrkA TM dimer (yellow) dissociated and assembled again without insertion of the protein, which interacted with water beads and the polar head beads (blue and green) of the lipids, as shown in the snapshot after 5 μs on the right.

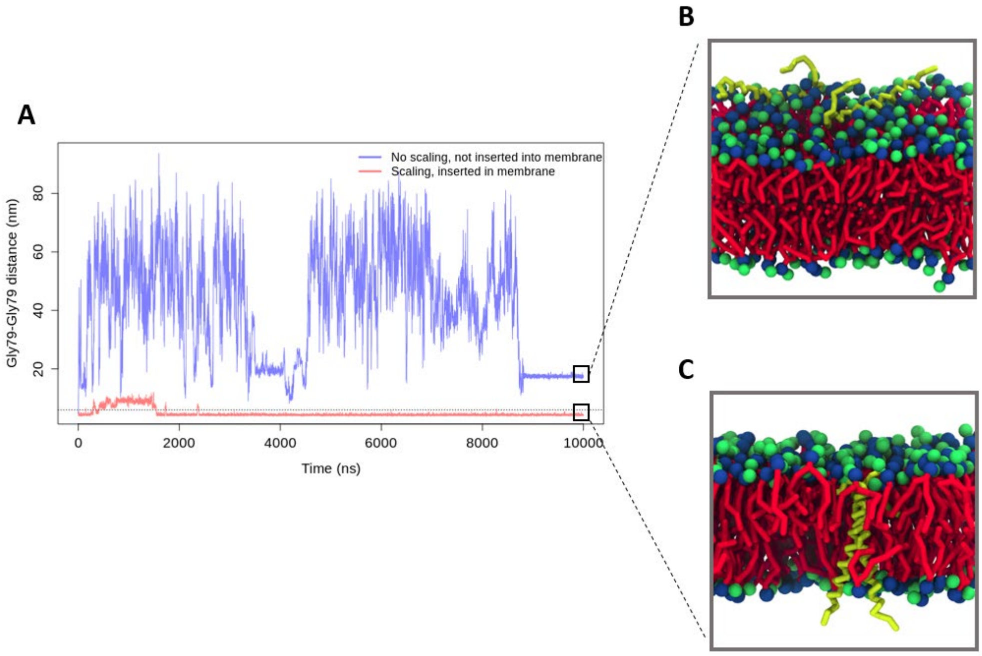


**Supplementary Figure 2: GpA monomers that are not embedded into the lipid bilayer tend to dissociate.** **A**: Distance between the central G79 residues of the two GpA monomers during two 10 μs simulations. In simulations without protein-water scaling and with no initial insertion of the GpA dimer into the membrane, high distances between the G79 residues indicate monomer dissociation (blue line). **B**: At the end of this trajectory, the distance between the two monomers stabilizes due to interactions between the protein and the polar head groups in the top bilayer leaflet. **C**: In contrast, the distance between the G79 residues of the two GpA monomers is below the contact cut-off distance (6Å, marked by the horizontal dashed line) during almost the entire trajectory when the protein dimer is first inserted into the lipid bilayer and then protein-water scaling is applied during the simulation (red line in **A**). The TM GpA dimer at the end of the simulation is shown in **C**.

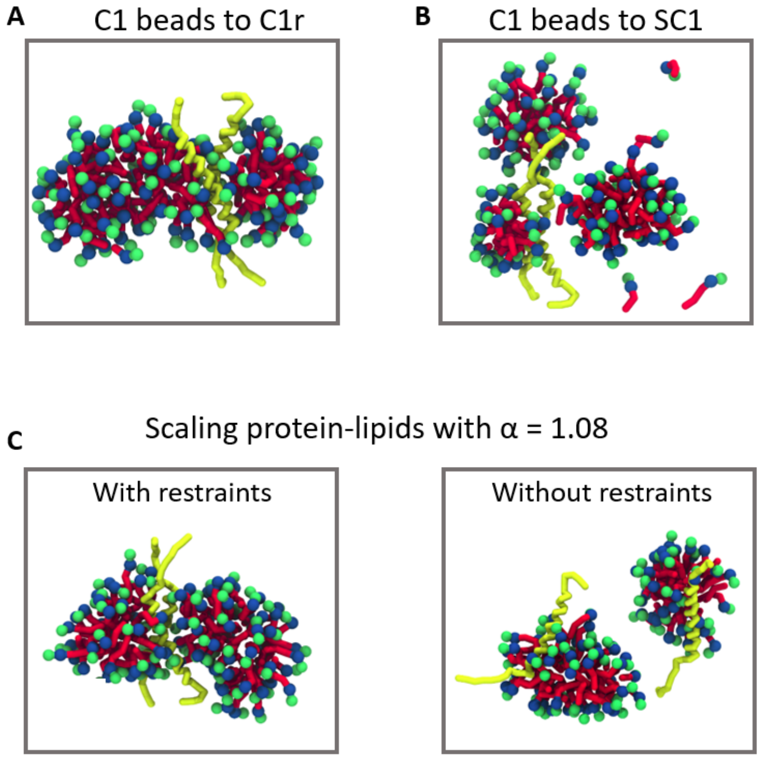


**Supplementary Figure 3: Different unsuccessful approaches to increase protein-lipid interactions and achieve insertion of the TrkA dimer into micelles. A**: Self-interactions between DPC tails were reduced by changing their constitutive C1 bead types to C1r. This label reduces self-interactions by increasing the interaction level between beads of the same type by 1. Insertion of TM TrkA dimers into micelles was not achieved with this approach. **B**: The size of the beads used in the DPC hydrophobic tails was reduced by changing them from C1 to SC1. Using this strategy not only did not result in protein insertion into micelles, but also resulted in a decrease of the lipid aggregation number. Not all DPC molecules participated in micelle assembly. **C**: A uniform scaling factor of $\alpha$=1.08 was introduced to increase non-bonded interactions between protein and hydrophobic C1 beads by 8%. Although a slight increase of protein-lipid interactions was observed, this was not sufficient for full insertion of the TM dimer into DPC micelles (left panel). Moreover, both monomers dissociated after release of the position restraints on the protein, ending in different micelles (right panel). Other values of the scaling factor from $\alpha$=1.04 to 1.16 were tested, yielding similar results (data not shown).

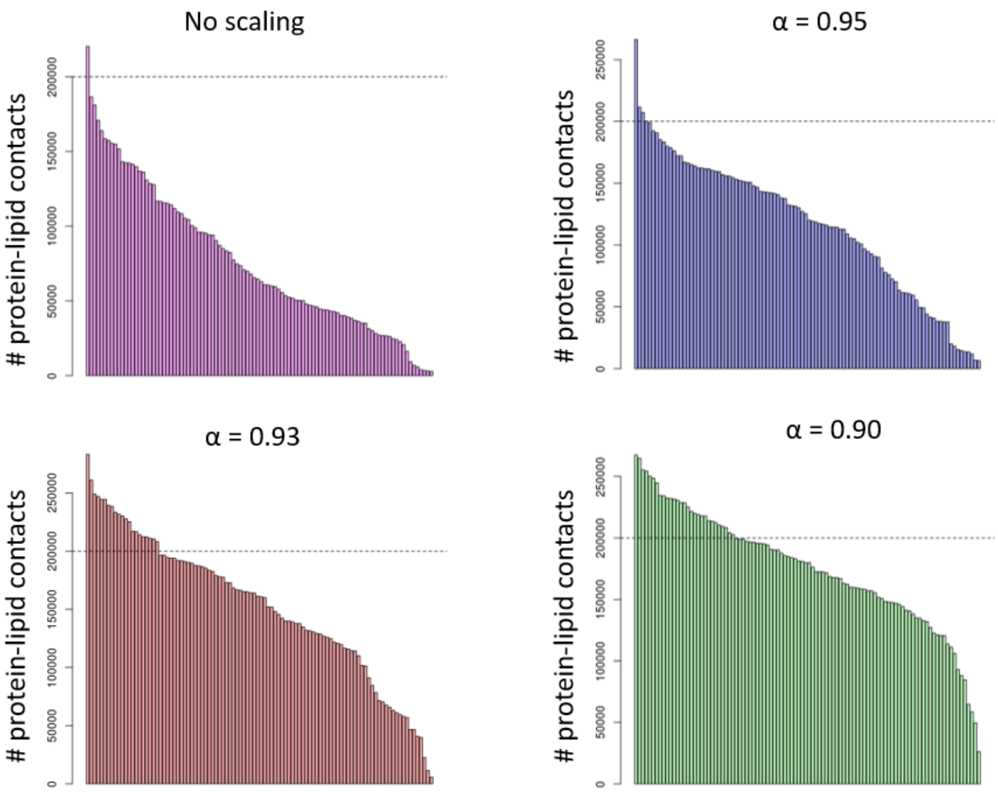


**Supplementary Figure 4: Number of contacts between GpA and POPC in simulations with different values of the scaling factor for protein-water non-bonded interactions.** 100 replicas of 60 ns duration were run with each scaling factor. Each bar shows the number of GpA-POPC contacts in one replica, with the replicas sorted in decreasing order. A threshold of 200,000 lipid-protein contacts (dashed line) is shown to identify insertion of GpA into the bilayer. Decreasing the scaling factor α led to higher numbers of replicas above this threshold: 1 for α = 1, 4 for α = 0.95, 20 for α = 0.93 and 30 for α = 0.90.

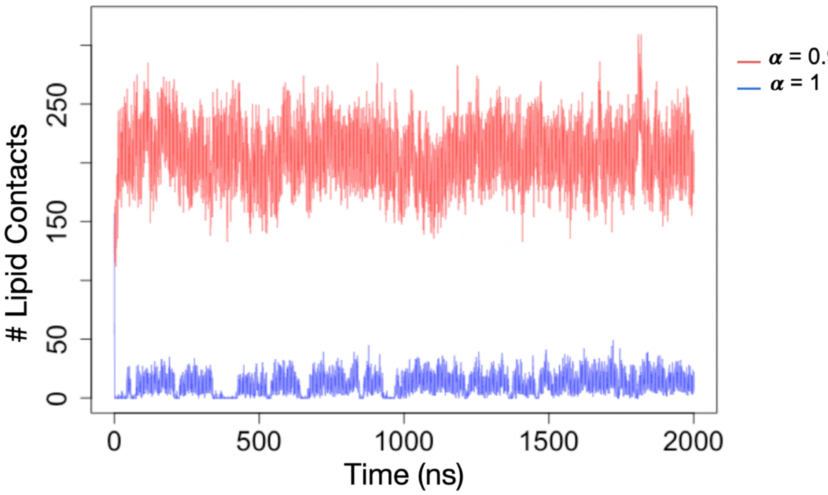


**Supplementary Figure 5: Number of contacts between GpA and the hydrophobic tails of the DPPC lipids during self-assembly simulations with and without scaling of the protein-water interactions in Martini 3.** The results of one replica simulation with (α = 0.9) and one replica simulation without (α = 1) scaling of the protein-water interactions are shown. Of the 10 replicas run for each system, similar behaviour was observed in 8 of the replicas with scaling and 9 of the replicas without scaling of the protein-water interactions. Thus, the GpA TM dimers encapsulate into the membrane upon DPPC bilayer self-assembly in most cases when the protein-water interactions are scaled by 10% and only rarely when the original unscaled Martini 3 parameters are used.

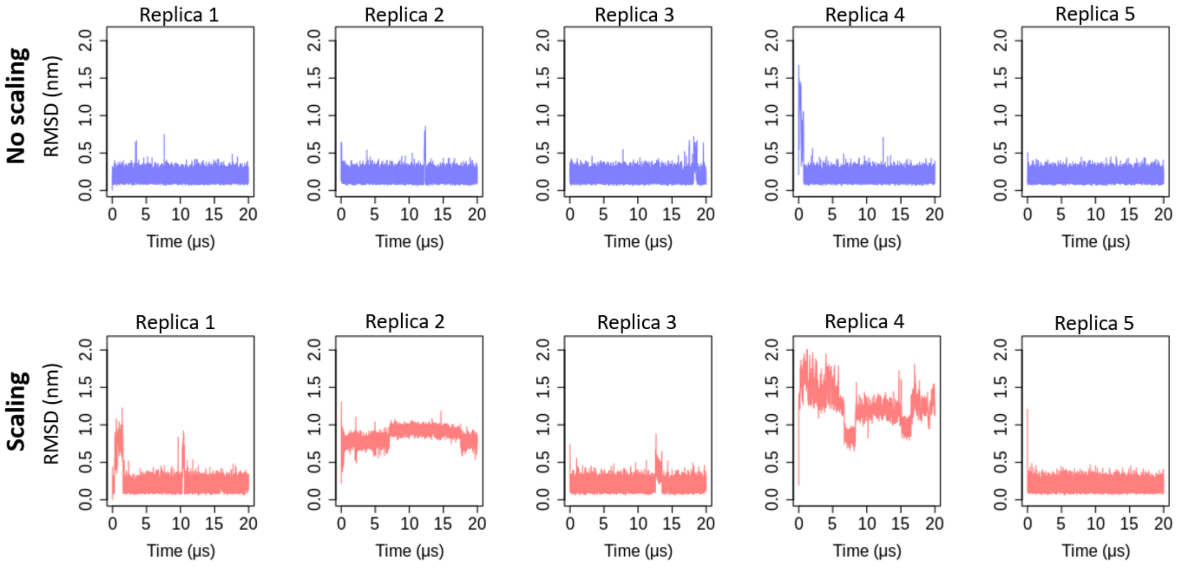


**Supplementary Figure 6: Root mean square deviation (RMSD) of the GpA dimer in different replicas with and without application of the scaling factor of α = 0.9 to reduce protein-water non-bonded interactions.** The NMR structure (PDB id 1AFO) was used as the reference structure.

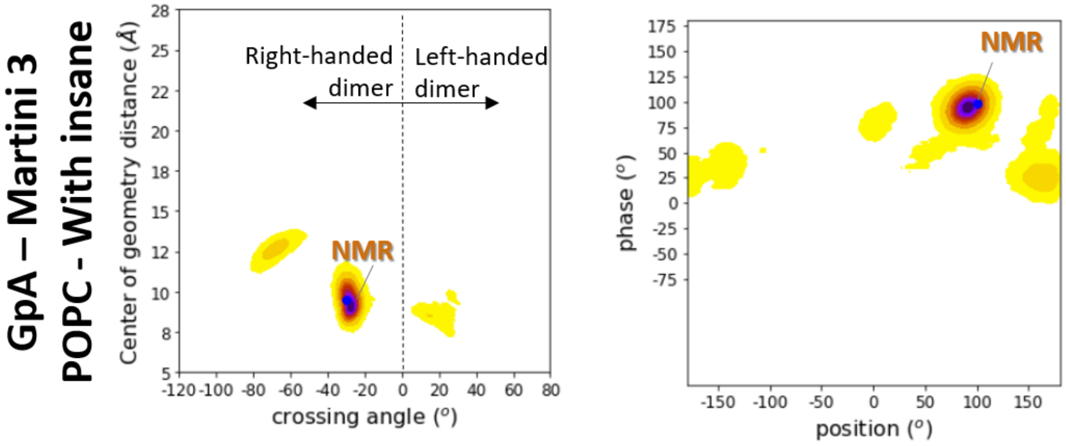


**Supplementary figure 7: Conformational landscape of GpA TM helix dimers in POPC bilayers inserted in the membrane via the *INSANE* method and simulated without introduction of a scaling factor using Martini 3.**

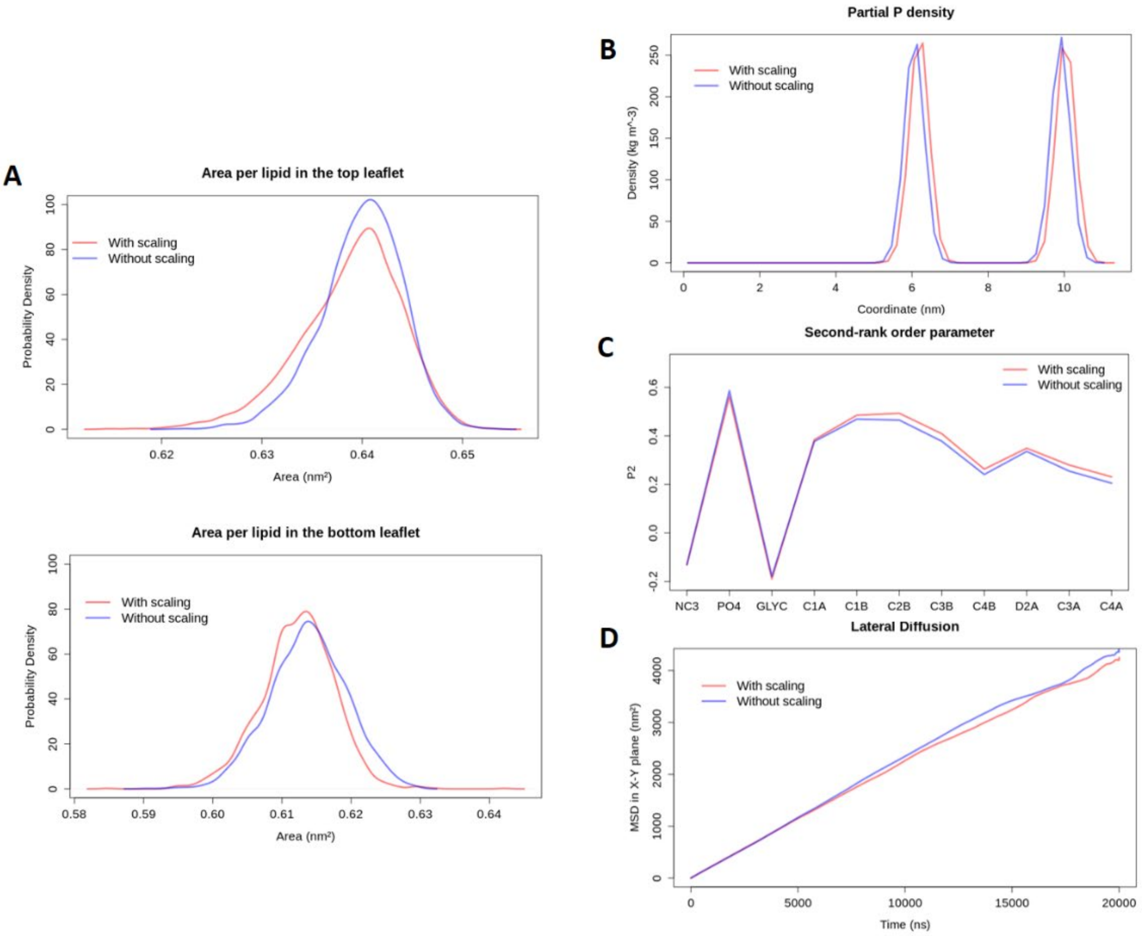


**Supplementary Figure 8: Structural properties of the POPC phospholipid bilayers do not differ in simulations using Martini 3 with and without application of the scaling factor of α = 0.9 to reduce protein-water non-bonded interactions. A**: Area per lipid in the upper and lower bilayer leaflets in simulations with and without scaling. Differences between the two leaflets indicate an asymmetric distribution of the lipids. **B**: Density of phosphate beads along the z-axis, which is orthogonal to the membrane plane. The distance between the two peaks gives an estimate of bilayer thickness. **C**: Second rank order parameter (P2) for consecutive bonds in POPC. **D**: Mean square displacement of the lipids in the XY plane as a function of time. The values are averaged over the entire trajectory. All plots were obtained from the same two replica trajectories (one with and one without scaling of protein-water interactions); the same behaviour was observed for all replicas (data not shown).

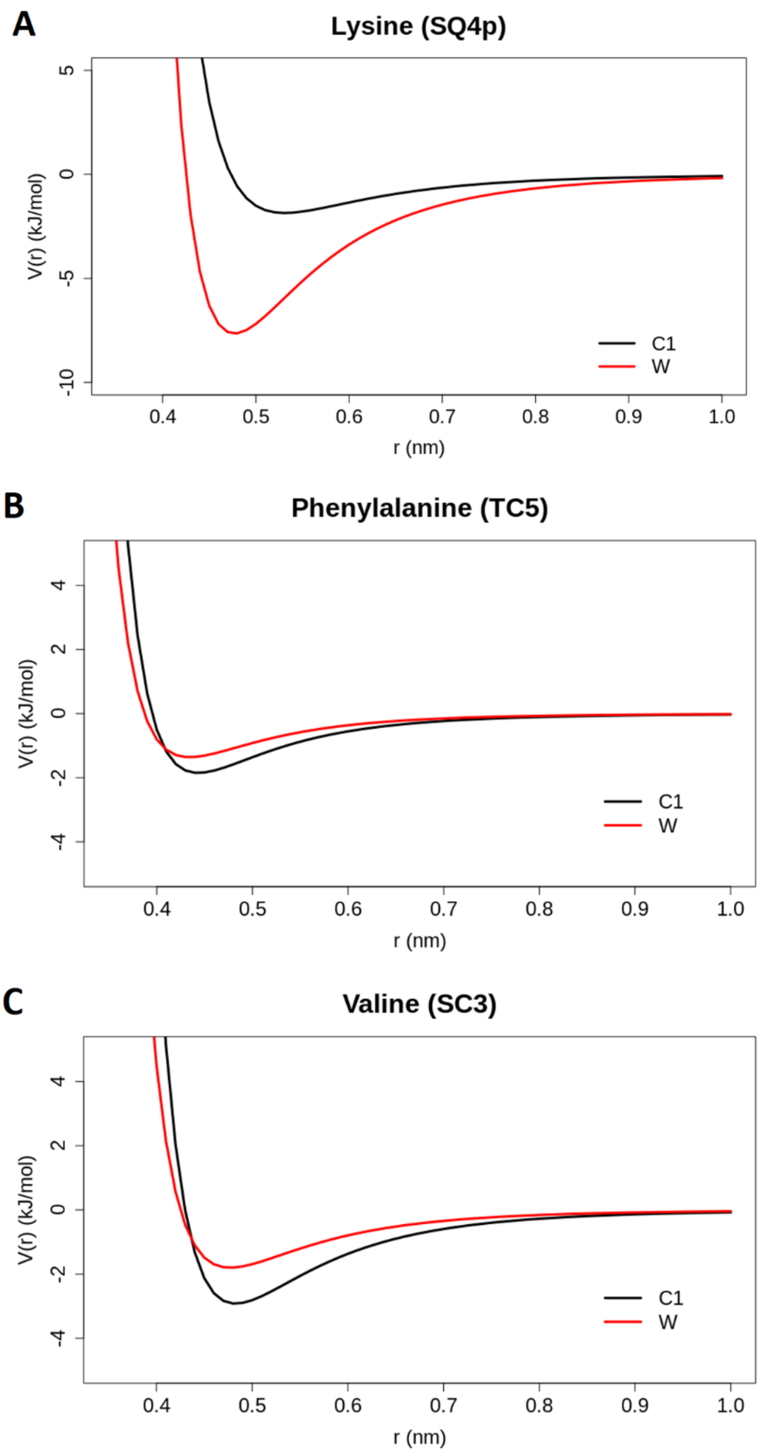


**Supplementary Figure 9: Lennard-Jones potentials of beads of selected residues with hydrophobic (black:C1) and water (red:W) beads in Martini 3**. **A**: Non-bonded interactions between the outermost side chain bead of lysine (SQ4p) and water beads are modeled by a Lennard-Jones potential with a large energy well depth. This reflects the hydrophilic nature of this part of this amino acid. **B**: The energy well depth of the Lennard-Jones potential between phenylalanine TC5 beads and hydrophobic beads of type C1 is just slightly greater than for interactions between TC5 and water beads. **C**: The Lennard-Jones potential between valine side chain beads (SC3) and C1 beads has a deeper energy well than that for SC3-W interactions, which reflects the hydrophobicity of this amino acid. To generate these plots, epsilon and sigma values of the given non-bonded interactions were obtained from the publicly available Martini 3 parameter file[1].

[1] http://cgmartini.nl/images/martini_v300.zip
